# Understanding the Pathogenicity of Parkin Catalytic Domain Mutants

**DOI:** 10.1101/2024.07.29.605699

**Authors:** Julian P. Wagner, Véronique Sauvé, Kalle Gehring

## Abstract

Mutations in the E3 ubiquitin ligase parkin cause a familial form of Parkinson’s disease (PD). Parkin and the mitochondrial kinase PINK1 assure quality control of mitochondria through selective autophagy of mitochondria (mitophagy). Whereas numerous parkin mutations have been functionally characterized and their structural basis revealed, several pathogenic PD mutations found in the catalytic RING2 domain remain poorly understood. Here, we characterize two pathogenic RING2 mutants, T415N and P437L and shed light on the underlying structural causes. For this purpose, we use biochemical *in vitro* assays in combination with AlphaFold modeling. We demonstrate that both mutants exhibit impaired activity using autoubiquitination and ubiquitin vinyl sulfone assays. After determining the parkin minimal ubiquitin binding region, we show that both mutants display impaired binding to the ubiquitin molecule charged onto the E2 enzyme. Finally, we employ the most recent version of AlphaFold 3 to generate a structural model of the phospho-parkin/phospho-ubiquitin/ubiquitin-charged E2 complex. This model consolidates our findings and provides a structural understanding for the pathogenicity of these two parkin variants. A better understanding of the different PD mutations at the molecular level can pave the way for personalized treatments and the design of small molecule therapeutics for the treatment of PD.

## Introduction

Parkinson’s disease (PD) is the second most common neurodegenerative disease and characterized by a set of clinical symptoms including motor symptoms, for instance tremors, rigidity and postural instability, and non-motor symptoms such as dysphagia or hyposmia. On a cellular level, a hallmark of PD is the loss of dopaminergic neurons in the *substantia nigra pars compacta*. Whereas the majority of cases are sporadic, 5-15% of patients present a familial form of the disease inherited through autosomal dominant or recessive mechanisms. A few highly penetrant mutations in different genes have been characterized, among them mutations in PINK1 and PRKN. Bi-allelic pathogenic PRKN variants are the most common forms of autosomal recessive early-onset PD (EOPD) (Kitada et al. 1998; Balestrino and Schapira 2020).

The gene PRKN encodes the E3 ubiqutin ligase parkin, a member of the RING-between-RING (RBR) family combining features of the RING-type and HECT-type families (Wenzel et al. 2011). Parkin is a multi-domain protein comprised of five domains (Ubl, RING0, RING1, IBR and RING2, sometimes referred to as Rcat) and two linker regions, one of them comprising the repressor element (REP). The catalytic core is formed by the four zinc-binding domains RING0, RING1, IBR and RING2 (Figure 1A). As for other members of the RBR family, the catalytic mechanism features binding of the E2 ubiquitin-conjugating enzyme to the RING1 domain, transfer of ubiquitin onto an active cysteine present on the catalytic RING2 domain via a transthiolation reaction and then onto a lysine residue of the substrate (Lechtenberg et al. 2016). Structural and biochemical studies over the last years have elucidated the activation process of parkin in detail, also providing insights into the functional effects of pathogenic PRKN mutations. Parkin activity in the cell is tightly regulated and under healthy cellular conditions, parkin is cytosolic and in an autoinhibited conformation: The catalytic cysteine is inaccessible due to intradomain contacts between RING2 and RING0 and the E2 binding site on RING1 is blocked by the UbI domain and the REP element (Trempe and Gehring 2023).

**Figure 1:**
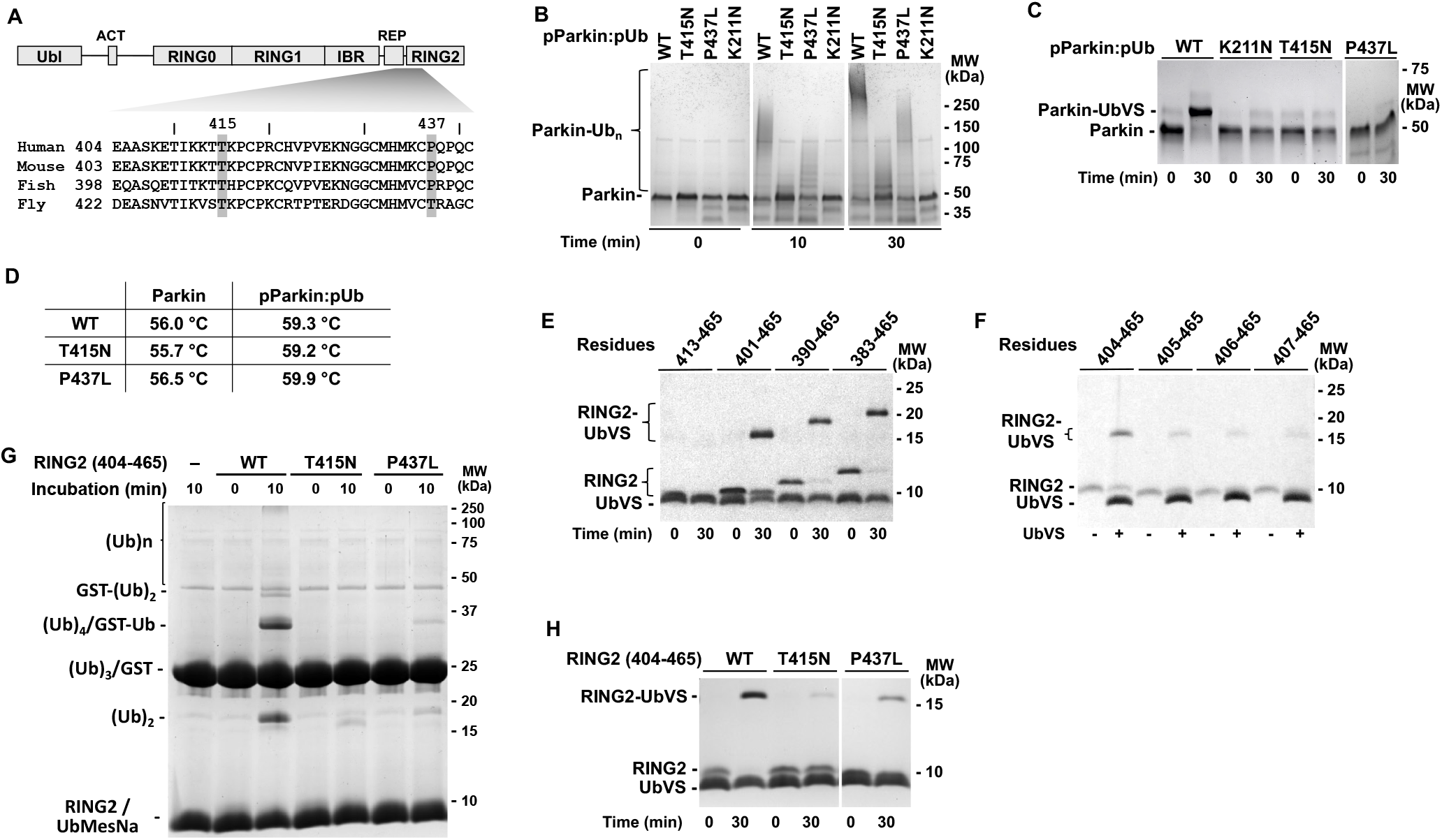
*In vitro* characterization of parkin mutants. (A) Parkin domain structure and sequence alignment of human, mouse, fish and fly parkin around the T415N and P427L mutations. (B) Autoubiquination assay of full-length activated parkin wild-type (WT), T415N or P437L. (C) UbVS assay of full-length wild-type (WT), T415N or P437L phosphorylated parkin in the presence of pUb. (D) Comparison of melting temperatures of inactive and activated wild-type (WT), T415N, and P437L parkin. (E-F) Determination of the minimal ubiquitin binding region using the UbVS assay with different RING2 constructs. (G) Bypass assay with wild-type (WT), T415N or P437L parkin (residues 404-465). (H) UbVS assays with wild-type (WT), T415N or P437L parkin (residues 404-465).

Together with the PTEN-induced kinase 1 (PINK1), parkin assures mitochondrial quality control through selective autophagy of damaged mitochondria (mitophagy). Mitochondrial damage leads to PINK1 dimerization, activation and stabilization on the surface of the mitochondria and to phosphorylation of ubiquitin (Ub). This in turn recruits parkin through binding to phosho-ubiquitin. Phosphorylation of the Ubl domain by PINK1 or alternatively, binding of a second phospho-ubiquitin induces large scale conformational changes (Fakih, Sauvé, and Gehring 2022). On the one hand, pUbI is released from RING1 thereby exposing the E2 binding site, on the other hand, pUbl binding to RING0 frees the catalytic RING2 domain. The catalytically active parkin now starts ubiquitinating substrates via the two-step mechanism described above. While it is established that loss-of-function mutations in either of the enzymes lead to a familial form of PD, a full picture of all the downstream consequences of the PINK1/parkin pathway is still missing.

The activation process has been studied in detail and atomic structures of inactive (Trempe et al. 2013), phospho-ubiquitin-bound (Kumar et al. 2017) and phosphorylated parkin (Gladkova et al. 2018; Sauvé et al. 2018) are available. However, less is known about how parkin interacts with the ubiquitin-charged E2 enzyme since no structure of the full complex formed by phospho-parkin/phospho-Ub and the E2∼Ub is available. Such a structure would allow to identify critical interactions required for thioester ubiquitin transfer. The currently available structure of UbcH7 bound to activated insect parkin lacks the charged ubiquitin, but it shows an interaction between the E2 enzyme and parkin RING1 through canonical binding sites, similar to other RBR E3 ligases (Sauvé et al. 2018). Structures of other RBR E3 ligases in complex with E2∼Ub have been published, including HOIP/UbcH5B and allosteric ubiquitin bound to HOIL-1 as well as the RNF216/Ub/E2∼Ub transthiolation complex. These complexes provided important insights into the distinct RBR catalytic mechanism and revealed a conserved transthiolation complex structure. In the complex, the E2 and E3 catalytic centres are ideally aligned for ubiquitin transfer and the RBR domain wraps around the ubiquitin loaded onto the E2 enzyme stabilising its open conformation and thereby preventing direct ubiquitin discharge. As also seen in the parkin structure lacking the RING2 and donor ubiquitin, the RING1 domain is the main E2 recruiting module. Other interactions between RING2/E2∼Ub are mainly mediated by ubiquitin itself, which contributes most of the binding surface, with only few direct interactions between the E2 enzyme and RING2. More specifically, the RBR RING1 binds E2 via the canonical RING1 central helix/E2 interface and the E2∼Ub isopeptide bond is placed into the RING2 active site primed for ubiquitin transfer. The allosteric ubiquitin is bound on the opposite side, i.e. binding the RING1-IBR domains (Wang et al. 2023). However, it is important to mention that despite these similarities and shared features, the different RBR family members have evolved specific features to exert their biological function highlighting once more the need for an active E2∼Ub/parkin transthiolation complex to provide us with a more thorough understanding of the parkin RBR E3 ligase. Furthermore, knowledge of the parkin structure in all its conformations and the structural basis for the pathogenicity of different variants is essential for structure-based drug design targeting this protein as a potential treatment for familial PD patients with PRKN mutations.

Here, we sought to understand the structural and functional effects of two parkin mutations T415N and P437L, which have been poorly understood until now. We characterized these variants biochemically and then determined the minimal ubiquitin binding region which we hypothesized to be the cause of their pathogenicity. We used the recently published version of AlphaFold 3 to predict the complete phospho-parkin/phospho-Ub/E2∼Ub complex and validated the model by binding and chemical crosslinking data. Thus, combining computational with biochemical data, we were able to provide the structural basis of the pathogenicity of these two parkin mutations.

## Results and Discussion

### Functional characterization of parkin mutants T415N and P437L and identification of the minimal functional RING2 domain

Functional and structural characterization of parkin enabled researchers to better understand and classify benign and pathogenic PD variants. However, despite these efforts, the structural basis for certain parkin variants remains unclear. Here, we focused on two single amino acid substitutions in the catalytic RING2 domain of the E3 ubiquitin ligase parkin, T415N and P437L (Figure 1A). Their mechanism of action remains poorly understood since the available parkin structures are insufficient to provide the structural basis for potential pathogenicity and explain their contribution to familial PD: Neither of them are located at positions that are obviously implicated in stability, zinc ion coordination or parkin catalytic activity. Previous *in cellulo* studies using the well-established mt-Keima reporter following mitochondrial damage (CCCP treatment) showed that both variants cause an impairment of mitophagy compared to WT parkin. Whereas P437L could be rescued by introduction *in cis* of two hyperactive variants (F146A and W403A), leading to an increase in mitophagy similar to the increase observed in the WT, T415N could not be rescued (Yi et al. 2019).

We first confirmed the impaired activity with auto-ubiquitination assays showing that T415N exhibits limited ubiquitination activity (Figure 1B). P437L also has a reduced ubiquitination activity, but not to the same extent. Next, we turned towards the ubiquitin vinyl sulfone (UbVS) assay: This assay uses a chemically reactive derivative that crosslinks to the parkin active–site cysteine (Borodovsky et al. 2001) and the formation of the UbVS adduct is therefore a suitable measure of exposure of the parkin catalytic site. This assay is widely used to evaluate parkin activation through RING2 exposure (Riley et al. 2013; Ordureau et al. 2014; Wauer et al. 2015) and is specific for ubiquitin. In controls with other ubiquitin-like VS reagents, e.g. Sumo-VS, no adducts are formed (data not shown). We compared the T415N and P437L variants with the WT and K211N, a known pathogenic variant featuring disturbed pUb binding during activation (Figure 1C). While the parkin-ubiquitin crosslinked band can be clearly seen for WT, it is absent for K211N, T415N and P437L. A caveat is that the UbVS assay cannot provide insight as to why the RING2 release is disturbed. On the one hand, it is possible that the active cysteine is not exposed because the RING2 domain is not released upon phosphorylation of their Ubl domain. This is the case for the K211N mutant. On the other hand, a loss of RING2 catalytic activity could also explain this observation.

To explore the first hypothesis, we evaluated thermal stability of parkin mutants in thermal shift assays to rule out the possibility that these mutants affect the stability of the protein. Compared to the WT, the mutants do not show any significant shifts in the melting temperature (T_m_) in both, non-activated and activated conditions (Figure 1D). This is in agreement with the mutants expressing similarly to WT in previous studies (Yi et al. 2019) as well as a recently published deep mutational scanning study evaluating the abundance of different parkin variants (Clausen et al. 2024).

Since the stability of the autoinhibited conformation did not appear to be affected, we investigated the second hypothesis (i.e. reduced activity of the parkin RING2) using UbVS assays. Starting with the RING2 domain plus the REP, we tested constructs with different N-termini to determine the minimal catalytically active parkin fragment. The longest fragment and deletions to residue 401 showed crosslinking while the fragment starting at residue 413 did not (Figure 1E). We repeated the experiment with single residue increments and determined parkin residues 404-465 to be the minimal fragment able to crosslink with UbVS (Figure 1F). This fragment was used in subsequent experiments.

### T415N and P437 impair the activity of the isolated RING2 domain

Having ascertained the minimal catalytically active region, we went on to examine reasons the T415N and P437L variants are impaired in their autoubiquitination activity. As the residues are not adjacent to the catalytic cysteine in RING2, we hypothesized that the mutations might interfere with the binding of ubiquitin. We thus used parkin constructs with residues 404-465 in a bypassing assay with a chemically activated ubiquitin C-terminal thioester probe, ubiquitin mercaptoethanesulfonate (UbMES), which mimics E2∼Ub (Park et al. 2017). The assay bypasses the necessity of E1 and E2 enzymes, since this compound directly reacts with catalytically active E3 enzymes. Using GST as a substrate and comparing to WT, we clearly see that the impaired activity must stem from impaired ubiquitin transfer from this minimal ubiquitin binding region since GST-ubiquitination is significantly lower for the mutants (Figure 1G). We repeated the UbVS assay with the same constructs to evaluate ubiquitin binding and saw significantly less crosslinking with the T415N and P437L variants compared to the WT (Figure 1H). In both experiments (bypass assay and UbVS assay), the P437L mutant showed less impairment than T415N. This is similar to the results in the autoubiquitination assays (Figure 1B). Taken together, these results demonstrate that T415N and P437L are functionally impaired due to inactivity of the RING2 domain, possibly due to a loss of affinity for ubiquitin binding.

### AlphaFold modeling of phospho-Ub:phospho-parkin E2∼Ub complex

To gain structural insights into the pathogenicity of the two variants, we used AlphaFold 3 (Abramson et al. 2024) to predict a computational model of the fully activated phospho-parkin/phospho-Ub/E2∼Ub complex. AlphaFold is an artificial neural network that predicts protein 3D structures based on amino acid sequence. The predictions are based on multiple sequence alignments and evolutionary constraints. With the most recent version of AlphaFold, post-translational modifications and metal ions can be included in the query and complexes between macromolecules can be predicted. We employed AlphaFold 3 using the sequence of human parkin with phosphorylation of the Ubl domain at serine 65, the sequence of human ubiquitin, of human phosphorylated ubiquitin at serine 65, the sequence of the E2 enzyme UbcH7 as well as eight zinc ions as query.

Using the standard protocol, five models were predicted; they were highly similar, suggesting a reliable prediction. Other quality indicators (ipTM and pTM scores > 0.7 and PAE plot) were good increasing our confidence in the model (Supplemental Figure S1). The overall conformation of the complex is similar to other published RBR structures (Wang et al. 2023) with the RBR module wrapping around the E2∼Ub and the allosteric phospho-ubiquitin bound on the opposite side interacting with the RING1 and IBR domains (Figure 2A). Importantly, the structure of the complex was consistent with the biochemistry of parkin activity. In the predicted model, the cysteine 86 of UbcH7 and the catalytic cysteine of parkin at position 431 are in close vicinity, primed for the transthiolation reaction. The C-terminal glycine of ubiquitin was placed in a way that would allow its transfer onto parkin (Figure 2B). In all five models, the orientations of amino acid side chains were consistent with the known mechanism of transthiolation, highlighting the robustness of the prediction. As also observed previously, interactions between E2∼Ub and the RING2 domain are mostly mediated through the ubiquitin charged on the E2 enzyme. An alpha helix formed by the REP domain and the N-terminal residues of the RING2 domain interacts with the hydrophobic surface on ubiquitin centered around isoleucine at position 44 (Figure 2C) to generate the open E2∼Ub conformation that favors transthiolation (Dove et al. 2016). Intriguingly, the helix includes W403, which promotes parkin derepression when mutated to alanine (W403A)(Trempe et al. 2013). In the AlphaFold model, the tryptophan interacts with F45 of ubiquitin, which would suggest that the W403A mutant should bind E2∼Ub less well. The hydrophobic contact provided by the tryptophan is apparently unnecessary as UbVS experiments show the RING2 domain is fully active without it (Figure 1F). While not part of the helix, the mutated residues T415 and P437 are adjacent and form hydrophobic contacts with ubiquitin to promote binding.

**Figure 2:**
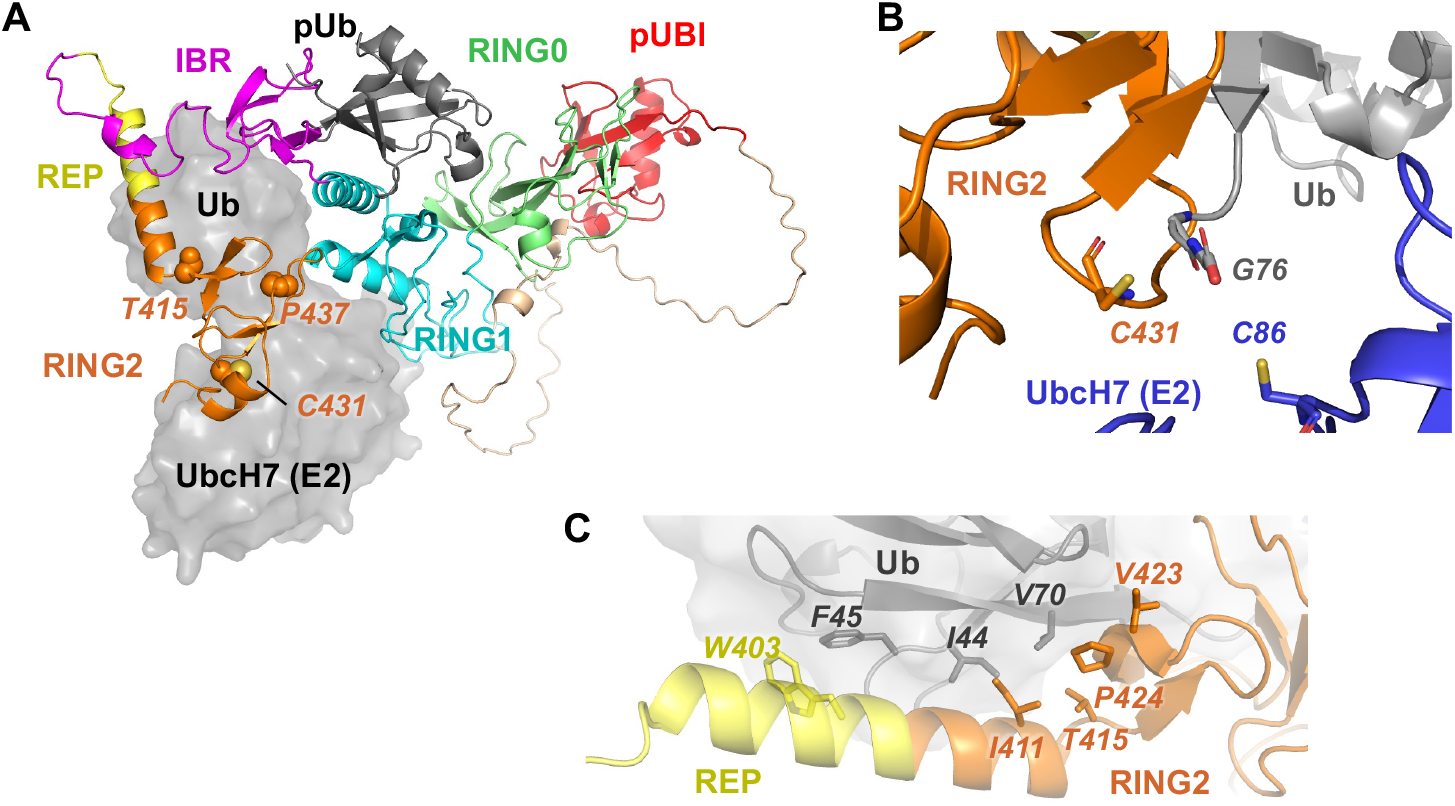
AlphaFold 3 model of the parkin complex with a ubiquitin-charged E2. (A) Overall conformation of the predicted model of phosphorylated parkin with bound allosteric phospho-ubiquitin and ubiquitin-charged UbcH7. A helix formed by the REP and N-terminus of RING2 positions the parkin catalytic site above the ubiquitin-E2 thioester. Residues 415 and 437 bind the ubiquitin on the E2. (B) Zoom on the catalytic centers primed for the transthiolation reaction. The catalytic cysteines on UbcH7 and on the RING2 domain as well as the C-terminal glycine of the ubiquitin are shown in stick representation. (C) Zoom on the ubiquitin interacting helix formed by residues from the REP and N-terminal portion of RING2. The interaction interface is largely hydrophobic centered around ubiquitin residue 44.

### Probing the ubiquitin/parkin interface and experimental validation of the model

We next aimed to experimentally validate the predicted model and assess the importance of the mutated residues for binding ubiquitin. As a first approach, we employed a *p*-benzoyl-L-phenylalanine (BPA) crosslinking assay (Chin et al. 2002). This non-canonical amino acid can be incorporated into the protein at a given position and upon irradiation with UV-A light, highly reactive short-lived species are formed capturing proximal molecules – within a radius of a few Å – with covalent bonds. On reducing SDS-PAGE, this will lead to a shift in the molecular weight and identification of crosslinked complexes. We modified six positions with the crosslinking reagent and examined whether a crosslink could be detected. Crosslinking was detected for residues T9, T66 and L71 (Figure 3A) which are closest to parkin residues in the AlphaFold model. Attempts to determine the parkin residues crosslinked by mass spectrometry were unsuccessful. The three other sites showed no crosslinking consistent with their greater distance from parkin residues (Figure 3B).

**Figure 3:**
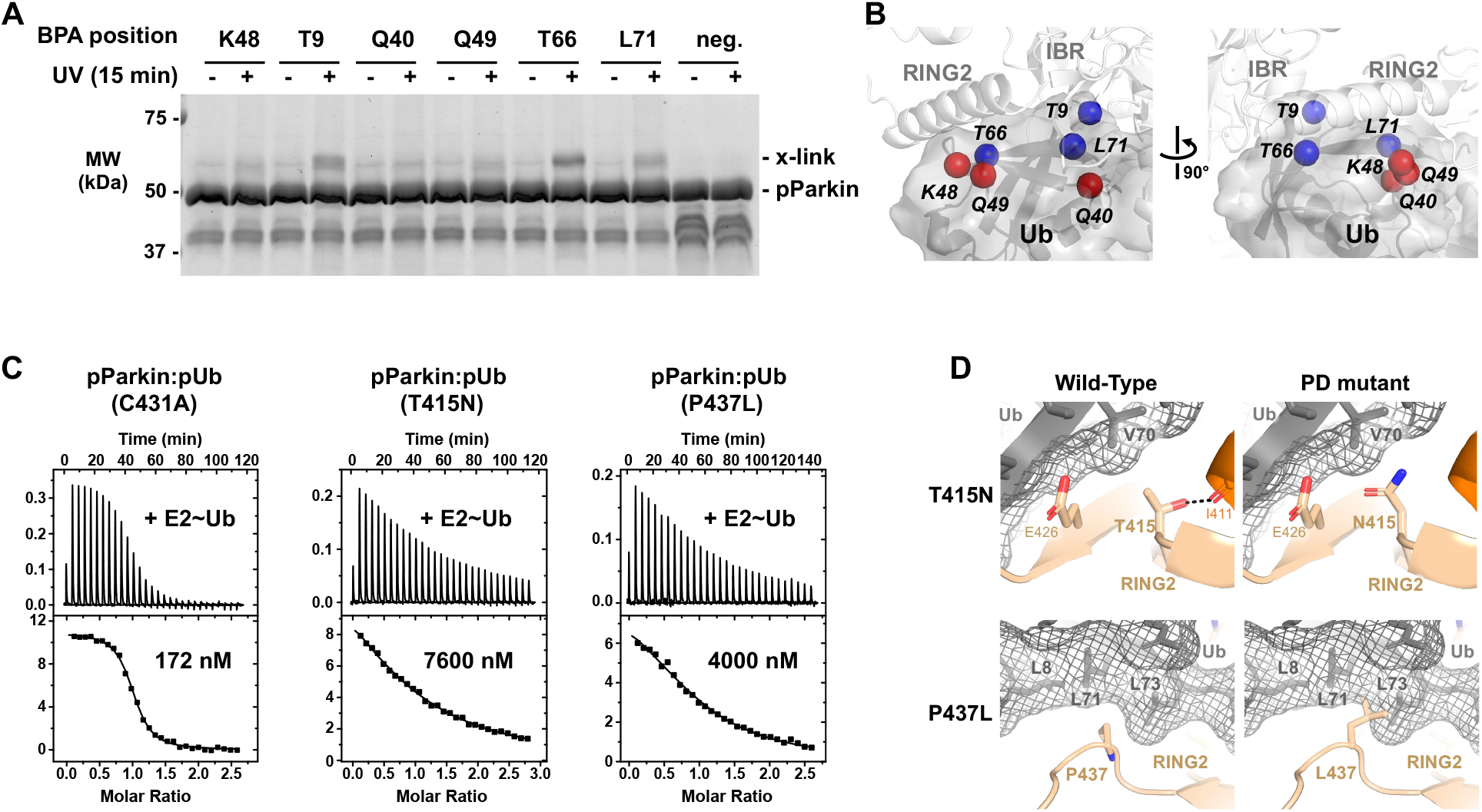
Structural basis for the pathogenicity of the parkin mutants. (A) Location of the six positions in ubiquitin modified with a *p*-benzoyl-L-phenylalanine (BPA) crosslinking reagent. Positions that generated crosslinks are colored blue; those that did not are colored red. (B) SDS-PAGE analysis of BPA crosslinking assay. Crosslinks were detected for BPA incorporated at residues 9, 66 and 71. Ubiquitin without BPA reagent was used as a negative control. (C) Isothermal titration calorimetry (ITC) experiments with phosphorylated parkin with pUb titrated with ubiquitin-charged E2 enzyme. The wild-type experiment used a catalytically inactive mutant (C431A) to prevent transthiolation and discharging of the E2 enzyme. (D) Zoom on residues T415 and P437 from the AlphaFold model showing loss of favorable binding contacts with ubiquitin.

In a second approach, we carried out ITC experiments with activated parkin to measure the affinity of UbcH7∼Ub binding (Figure 3C). In agreement with previous measurements (Kumar et al. 2015), the complex of phosphorylated parkin (pParkin) and pUb bind UbcH7∼Ub with submicromolar affinity. These experiments require the use of a catalytically inactive parkin mutant to prevents ubiquitin discharging during the titration. Titrations with both T415N and P437L parkin mutants showed a drastic decrease in K_D_ (∼40-fold and ∼20-fold, respectively). This confirms that both mutants are inactive due to a loss in affinity for ubiquitin, both when charged on the E2 enzyme in autoubiquitination reactions and when it is alone in the UbVS or bypassing reactions.

We used the computational model of the activated complex to further understand the molecular mechanism underlying the pathogenicity of the parkin mutants (Figure 3D). In a model of the T415N mutant, the hydrophilic side chain of asparagine is close to the ubiquitin hydrophobic patch and valine 70. This likely leads to repulsion and steric hindrance and explains the impaired ubiquitin binding. The AlphaFold model also provided insight into P437L mutant: the isobutyl sidechain of leucine is larger than the proline side chain, which leads to a steric clash with leucines 70 and 71 of ubiquitin.

In conclusion, we have used a variety of biochemical assays to show that parkin EOPD mutations, T415N and P437L, are impaired in transthiolation due to their inability to bind ubiquitin. The REP helix, which contributes to parkin inhibition in the autoinhibited structure, extends in the complex of activated parkin to include the first N-terminal residues of RING2. This helix binds ubiquitin to generate the open E2∼Ub conformation and promote ubiquitin transfer to the parkin catalytic cysteine. These insights into the structure of the fully activated parkin in complex with a ubiquitin-charged E2 enzyme will be a valuable resource for future studies and brings us one step closer to finding an effective treatment for patients.

## Experimental procedures

### Cloning, expression, and purification of recombinant proteins in *E. coli*

Single-point mutations and deletions were generated using PCR mutagenesis (Agilent) and proteins expressed in BL21 (DE3) *E. coli*. Purification of full-length and truncated parkin, UbcH7, Tc-PINK1 and human His-E1 were done using methods previously described (Sauve et al. 2015; Berndsen and Wolberger 2011). Briefly, proteins were purified by glutathione-Sepharose (Cytiva) or Ni-NTA agarose (Qiagen) affinity chromatography, followed by either 3C protease cleavage to remove the GST tag or Ulp protease cleavage to remove the His-SUMO tag. Size-exclusion chromatography was used as a last step. Phosphorylated Ub was produced and purified according to published procedures (Ordureau et al. 2014; Wauer et al. 2015). Purified proteins were verified using SDS-PAGE analysis. Protein concentrations were determined using UV absorbance.

### Autoubiquitination assays

The autoubiquitination assays of full-length WT and mutated parkin were performed at 22 °C for 10 or 30 min by adding 2 µM full-length phosphorylated WT, T415N or P437L parkin in complex with pUb, to 100 nM human His-Ube1, 2 µM UbcH7, 75 µM ubiquitin in 50 mM Tris pH 8.0, 150 mM NaCl, 1 mM TCEP, 5 mM ATP and 10 mM MgCl_2_. Reactions were stopped by the addition of reducing SDS-PAGE loading buffer and the level of ubiquitination was analyzed on SDS-PAGE gels stained with Coomassie blue.

### Differential scanning fluorimetry

To assess the thermal stability of the parkin mutants, 5 µM of non-phosphorylated or phosphorylated parkin in complex with pUb were mixed with 1X Protein Thermal Shift™ Dye kit (Applied Biosystems, Life Technologies) in 30 mM Tris, 150 mM NaCl, 1 mM TCEP, pH 8.0. The samples were heated from 25 °C to 99 °C in 1% steps using StepOnePlus (Applied Biosystems, Life Technologies). Data were analyzed using Thermal Shift software (Life Technologies). The maximum change of fluorescence with respect to temperature was used to determine the denaturation temperature. Experiments for each sample were performed in triplicate.

### Ubiquitin vinyl sulfone assays

For ubiquitin vinyl sulfone assays, 2 µM parkin (either full-length or truncated parkin) were incubated in the presence of 10 µM UbVS (R&D Systems) in 30 mM Tris-HCl pH 8.0, 150 mM NaCl, 1 mM TCEP, pH 8.0 for 30 min at 37 °C. Reactions were stopped by the addition of reducing SDS–PAGE loading buffer and the level of crosslinking was analyzed on SDS–PAGE gels (Bio-Rad) stained with Coomassie blue.

### Bypassing assay

The bypassing assay was adapted from (Park et al. 2017). Ubiquitin-MESNa was prepared by incubation of 100 µM ubiquitin, 250 nM His-E1, 100 mM MesNa, 10 mM MgCl_2_ and 10 mM ATP at 37 °C for 5 h in 0.1 M NaPO_4_, pH 8.0. The reaction was stopped by dialysis against 20 mM HEPES, 120 mM NaCl, pH 6.5. The ubiquitin-MESNa was further purified by gel filtration in 20 mM HEPES, 120 mM NaCl, 0,5 mM TCEP, pH 7.4. The bypassing assays of WT and mutated REP-RING2 (residues 404-465) were performed at 22 °C for 10 min by adding 1 µM WT, T415N or P437L REP-RING2, to 30 µM ubiquitin-MESNa and 30 µM GST protein in 30 mM HEPES pH 8.0, 150 mM NaCl, 0.5 mM TCEP. Reactions were stopped by the addition of reducing SDS-PAGE loading buffer and the level of ubiquitination of GST was analyzed on SDS-PAGE gels stained with Coomassie blue.

### AlphaFold modeling

The structure of the phospho-ubiquitin bound phospho-parkin in complex with ubiquitin-charged E2 enzyme was predicted using the AlphaFold server (Google DeepMind and Isomorphic Labs: https://alphafoldserver.com/, powered by AlphaFold 3) (Abramson et al. 2024). The query consisted of the amino acid sequence of human parkin with phosphorylated serine 65, a ubiquitin with phospho-serine at position 65 as well, human UbcH7 (E2 enzyme), an unmodified ubiquitin and eight zinc ions (Zn^2+^). The prediction was run using standard settings, i.e. five models were predicted and corresponding PAE plots produced. Models were visualized in PyMOL and PAE plots were visualized using a custom python script.

### Isothermal titration calorimetry (ITC)

ITC measurements were carried out at 20 °C using VP-ITC (Microcal). Samples were in 50 mM Tris-HCl, 150 mM NaCl, 1 mM TCEP, pH 7.4. Proteins (10 µM WT, T415N or P437L pParkin:pUb) were titrated with one injection of 5 µl followed by 28 injections of 10 µl of 100 µM ubiquitin-charged UbcH7. Data were fitted to a single set of identical sites using Origin v7 software.

### *p*-Benzoyl-L-phenylalanine (BPA) crosslinking experiments

For expression of GST-tagged ubiquitin for the crosslinking experiments, the non-natural amino acid p-benzoyl-L-phenylalanine (BPA) was incorporated in response to the amber codon, TAG, in *E. coli*. The BPA-containing ubiquitin was then purified using glutathione-Sepharose (Cytiva) affinity chromatography followed by 3C protease cleavage to remove the GST tag. Subsequently, size-exclusion chromatography was carried out. Concentration was checked by UV absorbance and samples were sent to mass spectrometry to assess incorporation of BPA and degree of contamination by protein prematurely terminated due to incorporation of a STOP codon at TAG. To charge the E2 enzyme with the BPA-containing 100 nM human His-UBE1, 60 µM UbcH7 with 4 mM ATP and 8 mM MgCl_2_ were mixed and added to 200 µM Ub-BPA in 30 mM Tris-HCl pH 8.0, 150 mM NaCl, 1 mM TCEP. The mixture was incubated for 1 h at 37 °C. Successful charging of UbcH7 was checked by SDS-PAGE. Afterwards, phospho-parkin:pUb (C431A mutant to prevent ubiquitin discharging) was added to a final concentration of 5 µM. Samples were UV-radiated at 365 nm for 15 min at 4 °C or kept at 4 °C without irradiation as a negative control before SDS-PAGE analysis using reducing SDS-PAGE loading buffer with DTT.

## Data availability

All data described in this study are contained within the main article.

## Conflict of interest

The authors declare that they have no known competing financial interests or personal relationships that could have appeared to influence the work reported in this paper.

## Funding and additional information

This work was supported by Michael J. Fox Foundation grant (MJFF-019029) and Canadian Institutes of Health Research grant (FDN 159903).

## Notes

### Competing Interest Statement

The authors have declared no competing interest.

