## Supplemental Figure S1 for "Understanding the Pathogenicity of Parkin Catalytic Domain Mutants"

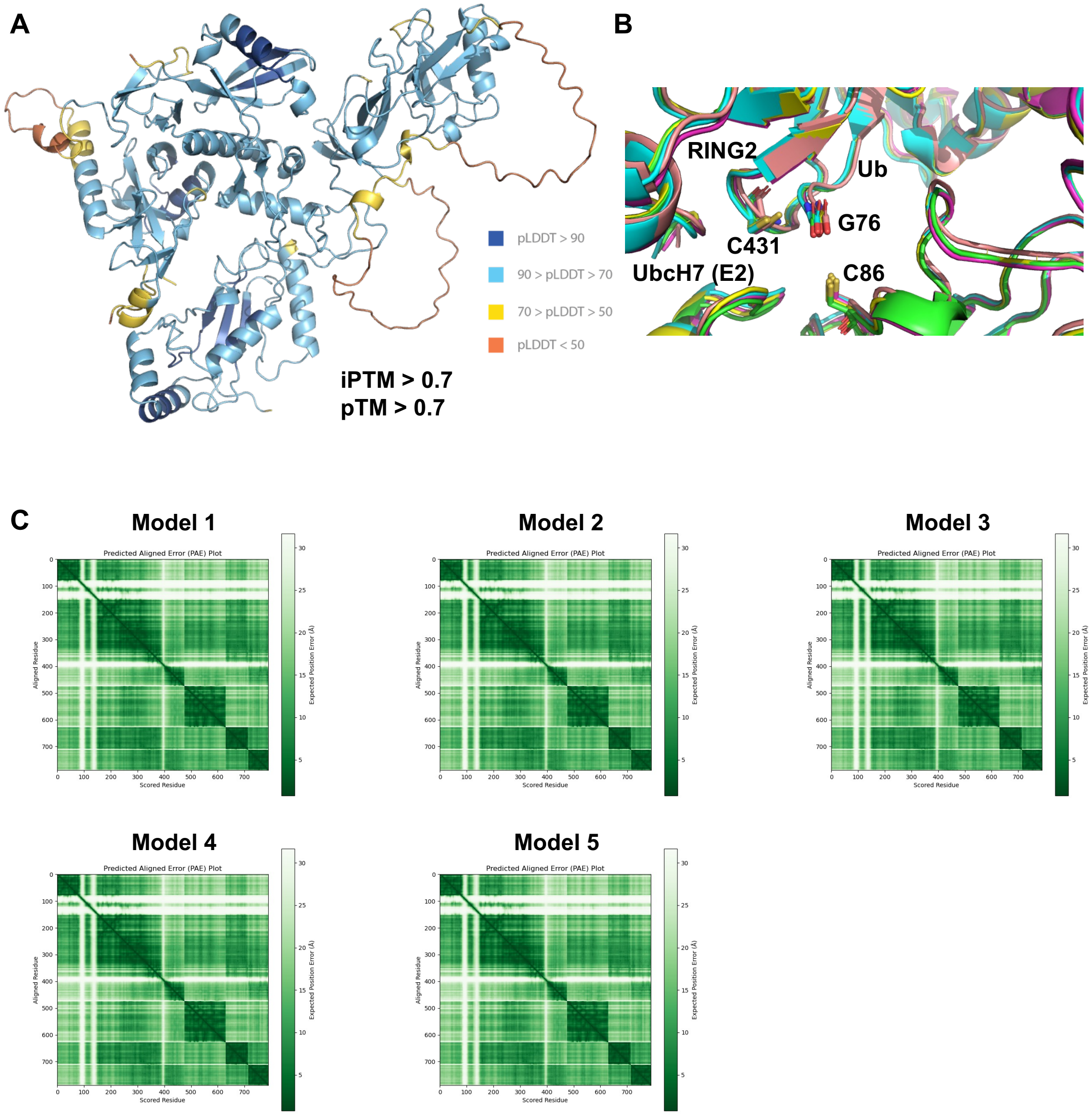


**Supplementary Figure 1:** AlphaFold 3 predicts the model of the complex with high confidence. (A) Model of phosphorylated parkin with bound allosteric phospho-ubiquitin and ubiquitin-charged UbcH7 colored according to pLDDT scores (local confidence). (B) Zoom on the overlay of the five models showing the parkin/E2 interface with the catalytic cysteine on UbcH7 and on the RING2 domain as well as the C-terminal glycine of the ubiquitin in stick representation. (C) Predicted aligned error (PAE) plots for the five predicted models. The PAE plot can be seen as a model for global confidence in the relative position of all pairs of residues.
